# Transfer of orientation memories in untethered wood ants (*Formica rufa*) from walking in an arena to walking on a motion compensation treadmill

**DOI:** 10.1101/2020.05.29.084905

**Authors:** Roman Goulard, Cornelia Buehlmann, Jeremy E. Niven, Paul Graham, Barbara Webb

**Author notes:** Corresponding author: Roman Goulard.

## Abstract

The scale of natural insect navigation during foraging makes it challenging to study, in a controlled way, the navigation processes that an insect brain can support. Virtual Reality and trackball setups have offered experimental control over visual environments while studying tethered insects, but potential limitations and confounds introduced by tethering motivates the development of alternative untethered solutions. In this paper we validate the use of a motion compensator (or ‘treadmill’) to study visually-driven behaviour of freely moving wood ants (*Formica rufa*). We show how this setup allows naturalistic walking behaviour and motivation over long timeframes. Furthermore, we show that ants are able to transfer associative and navigational memories from classical maze and arena contexts to our treadmill. Thus, we demonstrate the possibility to study navigational behaviour over ecologically relevant durations (and virtual distances) in precisely controlled environments, bridging the gap between natural and highly controlled laboratory experiments.

**Summary statement:** We have developed and validated a motion compensating treadmill for wood ants which opens new perspectives to study insect navigation behaviour in a fully controlled manner over ecologically relevant durations.

## 2 Introduction

Social insects often need to forage for sparsely distributed food sources in extended complex environments. To achieve this they utilise efficient navigation strategies, travelling long distances to find food to bring back to the nest [ants: (Collett and Collett, 2002; Cheng et al., 2014; Knaden and Graham, 2016), bees: (Menzel et al., 2005)]. Field studies and lab experiments have been used to investigate the underlying mechanisms of sensorimotor control and memory that drive such efficient navigation, but have inherent limitations. For instance, detailed observations of an animal’s path are only possible over short distances and for short durations, compared to natural foraging. Furthermore, identification of sensorimotor transformations, as for any reverse engineering method (Villaverde and Banga, 2014), needs precise control or observation of the coupling between the output (behaviour) and the input (sensory stimulation), which is difficult in the field. As an alternative, virtual reality (VR) methods provide a complementary tool (Schultheiss et al., 2017) to systematically control the access to sensory information for an individual while simultaneously recording precise behavioural responses, potentially over durations equivalent to long distance foraging.

One commonly used apparatus for establishing VR with walking insects is an air- or bearing-suspended trackball (Schultheiss et al., 2017) or wheels (Hedwig, 2017) driven by a tethered animal. However, tethering will inevitably induce physical discrepancies in the sensorimotor control of behaviour (Paulk et al., 2015), potentially impacting on the quantitative identification of sensorimotor transformation(s) from the analyses of input-output relationships. It may also affect the motivation of the animal (personal communications), thus limiting possible experimental durations. In particular, tethering often requires significant manipulation of the animal, which can induce qualitative changes in the drive to perform a particular task. It also reduces the ease of transferring animals between the real world and VR, e.g., for interleaved training and testing. Finally, a tether might directly constrain or prevent some aspects of the behaviour that we might wish to observe, such as the details of gaze control or posture, when a subject explores a visual scene. As a consequence, it is highly desirable to have a VR setup in which navigating insects can walk and behave freely.

In this paper, we adapt for use in wood ants (*Formica rufa*) a motion compensator setup, previously proposed to study freely moving beetles (Götz and Gambke, 1968), honeybees (Kramer, 1976), crickets (Weber et al., 1981) and spiders (Stowers et al., 2014). This type of setup has been more recently modified to study olfactory guided behaviour in moths (Shigaki et al., 2016), demonstrating its efficacy. The general approach of ‘motion compensation’ is to have the animal walking, untethered, on top of a servosphere, and to use a camera to detect its motion and then to move the sphere beneath the animal to compensate.

The latencies of early motion compensation systems, e.g. (Kramer, 1976), at around 150-400 ms, may have led to erroneous conclusions (Schmitz et al., 1982) and limited the popularity of this approach. However, current technology, such as high-speed cameras and efficient microcontrollers, enable the implementation of full compensatory closed loop control with high temporal resolution (Shigaki et al., 2016). Using a motion compensator system for robust control of insect position has mostly been used for analysis of taxis behaviours guided by sound (Weber et al., 1981; Schmitz et al., 1982), olfaction (Kramer, 1976; Sakuma, 2002; Shigaki et al., 2016) or vision (Stowers et al., 2014), but it has significant potential for the study of more complex navigation behaviours.

Our aim is to demonstrate that it is plausible to use a motion compensation treadmill to observe naturalistic behaviour of untethered navigating wood ants, to gain mechanistic insight into their visual navigation. Wood ants are a good model to study visual memory and navigation as they have been shown to produce reliable navigation in indoor laboratory experiments (Durier et al., 2003; Graham et al., 2003). We demonstrate memory transfer from traditional conditioning experiments and landmark navigation experiments to the treadmill apparatus. Furthermore, we show that disturbances created through the control of the servosphere (e.g. vibration or jitter) do not alter ant movement statistics, and we show, with trials of up to two hours, that ants remain motivated to walk in the conditions experienced on the treadmill. The results demonstrate the potential of the treadmill system for studying naturalistic orientation behaviours.

## 3 Materials & Methods

### 3.1 Motion compensation treadmill

The treadmill (Figure 1A) is based on a setup previously proposed to study moth olfactory responses (Shigaki et al., 2016). Ants walk on a white 3-DOF servosphere (foam, diameter: 120*mm*) supported by three rotors (stepper motors, SANYO SY42STH38 - 0406A) underneath, each one equipped with a dual-disc omniwheel (aluminium, diameter: 60*mm*, 2 discs, 5 rollers/disc) to avoid creating friction when the ball has to rotate perpendicularly to the axis of the rotor. Rotors are controlled using an Arduino Uno connected via USB to a computer (running Ubuntu 17) and three motor drivers (STMicroelectronics, ULN-2064B). The rotation of the ball is controlled by the ant’s motion which is tracked by a high-speed zenithal camera above the sphere. The camera is placed 150*mm* above the top of the sphere and we used a 12*mm* focal lens objective. The servosphere is lit from above by 4 lights, each consisting of 3 white LEDs and a diffusive cover to produce homogeneous lighting to facilitate tracking. A homogeneous cardboard ceiling, holding the LED lights, also blocks visual cues from above the setup. A white board, with a 5*cm* diameter hole to let only the top of the servosphere pass (figure 1D), covers the bottom of the setup, and a circular barrier of white flexible plastic closes the experimental chamber all around (figure 1A) and offers a support to set visual cues (figure 1D).

**Figure 1:**
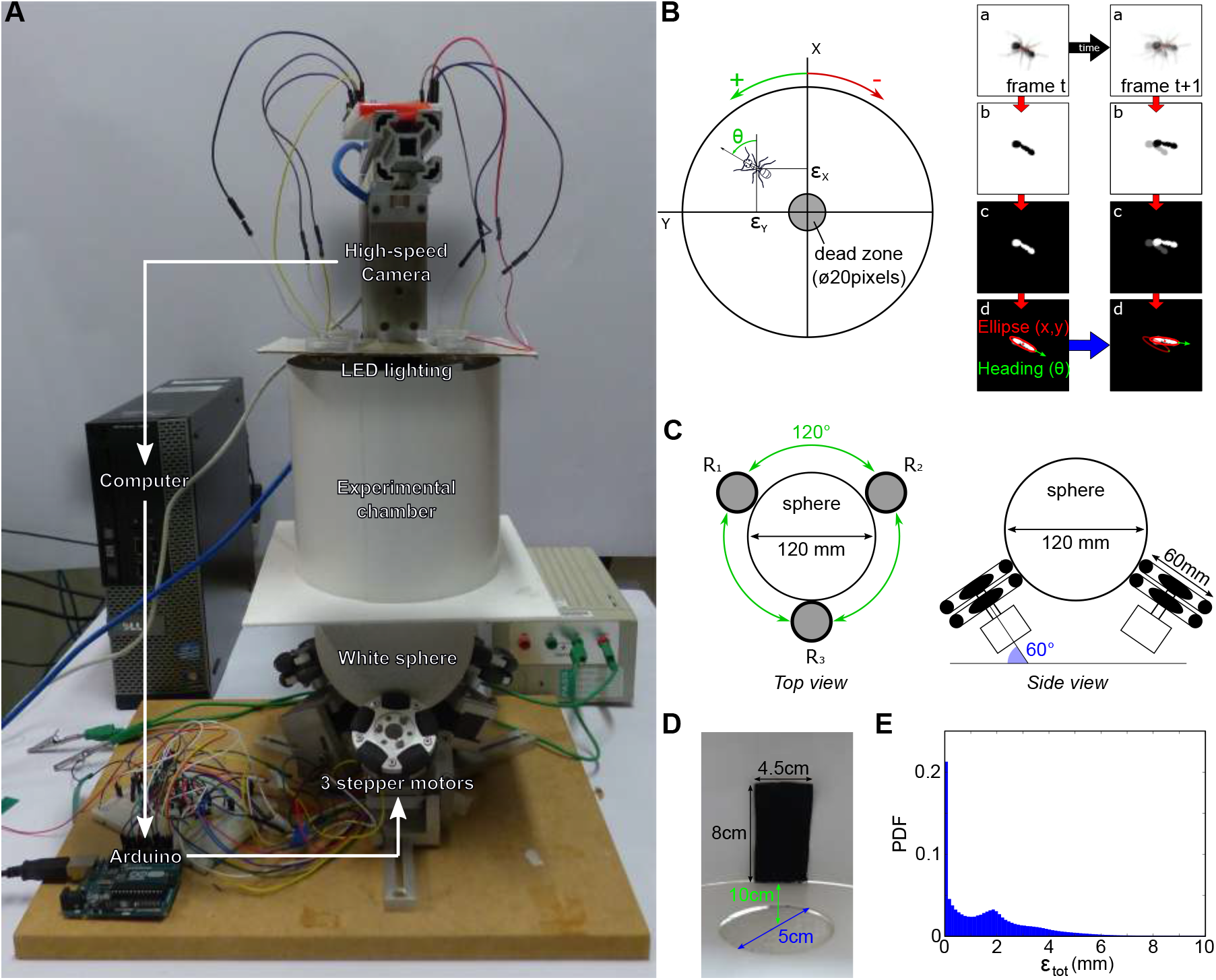
Treadmill for freely moving ants. (A) The motion compensator experimental set-up. (B) The tracking program extracts the position (*error X, error Y*) and orientation (*θ*) of the ant on the servosphere. The orientation is tracked with an uncertainty of ±180° which is corrected afterwards assuming ants mostly move forward. The left panels show the different stages of the tracking procedure: acquisition of an optically blurred image (a), binarization (b), contrast inversion (c) and ellipse detection (d). Tracking consistency between two consecutive frames is ensure by minimizing the centroid speed (Δ_*XY*_) and the heading change (Δ_*θ*_). Only the X and Y position are used to monitor the servosphere (see figure 2). (C) Top and side view from the system rotors-servosphere. The azimuth (120°) and the elevation (60°) angle are used to compute the rotational matrix used to transform the rotation of the sphere into the rotors reference frame (equation 4. (D) Inside view of the treadmill experimental chamber and of the piece of black fabric used as a visual cue during the experiments. (E) Probability density function of errors from the center measured by the tracking program along a batch of 174 experiments (10 to 15 minutes each).

#### 3.1.1 Tracking the ant

The position and 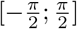 orientation of the ant are tracked using a high-speed camera (Basler ace acA640- 750um - monochrome, 700fps) and a custom python program. Ants are detected by binarizing the frame, using a threshold (set by the experimenter to lower background noise), and reversing the contrast, to make ants appear as a white blob on a black background (figure 1B). Then, contours are extracted from which we selected the blob most likely to be the ant, based on approximate size and minimization of the distance to the previously detected location. From the selected blob, we extract the position of the centroid and the 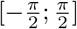 orientation by approximating an ellipse (facilitated by the blurring from the unfocused camera lens). The heading of the ant is approximated with a ±*π* uncertainty, which is resolved in post-processing, by assuming the ant moves forwards and kept consistent by minimizing the heading change between two consecutive frames (figure 1). As a consequence the ±*π* confusion persists when the ant is not moving so subsequent analysis involving orientation excludes periods of inactivity.

#### 3.1.2 Moving the 3-DOF servosphere

The X and Y distance from the center are used to actuate the rotation of the sphere and keep the ant at the center of the treadmill. In this work, we only controlled the position of the ant on the treadmill and let it freely rotate in a closed-loop manner. The extracted coordinates of the ant, 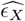 and 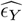 in pixels, are used to drive two independent Proportional-Derivative controllers to define the proper angular speed to apply to the ball 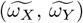. Both controllers are set with the same parameters and 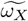 and 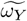 are calculated as follows:

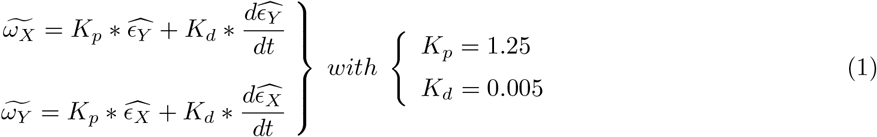

Then, *ω_X_* and *ω_Y_* are transformed into the reference frame of the three rotors’ to define the motor input necessary to actuate the sphere properly. The transformation matrix is computed using the azimuth and elevation of the rotors’ axes of rotation:

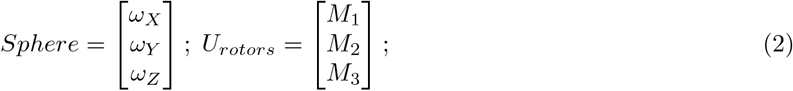

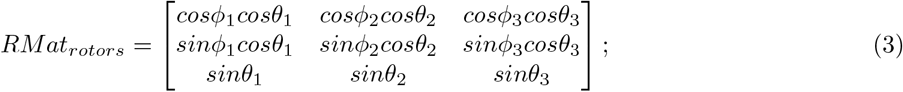

With *ω_X,Y,Z_* being the actual angular speed of the servosphere, *M*_1,2,3_ the motor speed of rotation, *ϕ*_1,2,3_ the elevation angle from the horizontal plane of the rotors’ axis of rotation and *θ*_1,2,3_ the azimuth angle, i.e., the orientation in the horizontal plane.

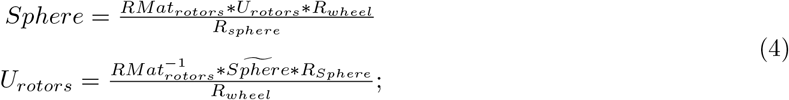

With *Sphere* being the actual dynamics of the sphere, 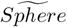 the desired dynamics of the sphere, *R_wheel_* the radius of the omniwheels and *R_sphere_* the radius of the sphere. From equation 4 it is determined what motor inputs will drive the sphere at the determined angular speeds (after the calculation of 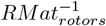, equation 6).

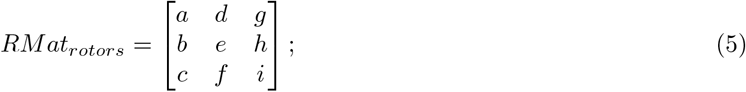

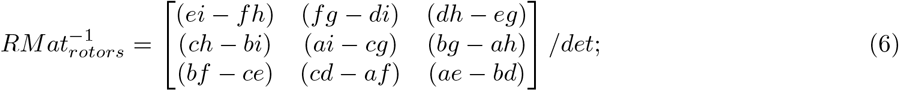

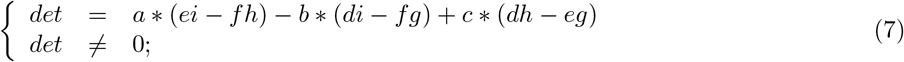

For the present setup, rotors are placed equally around the ball on the XY plane [*θ*_1_ = 0, *θ*_2_ = 120° and *θ*_1_ = −120°] and oriented around 60°from the horizontal [*ϕ*_1,2,3_ = 60°] (see figure 1C). The 3 rotor inputs are sent to the Arduino board via serial communication. The resulting feedback loop to control the treadmill is maintained at a frequency of around 500Hz. The full system is represented in the block diagram presented in figure 2.

**Figure 2:**
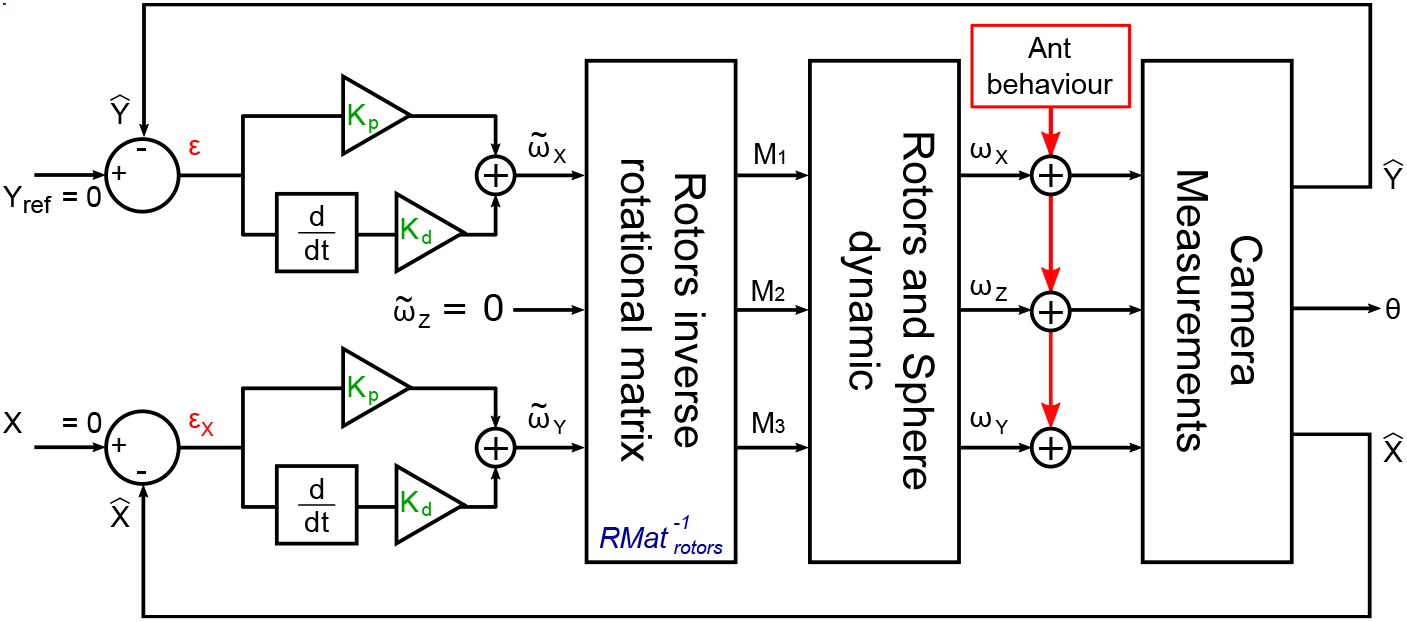
Block diagram of the complete servosphere Proportional-Derivative control. The coordinates of the ant are used to calculate the errors *ϵ_X_* and *ϵ_Y_* from the centre of the ball respectively along the X and Y axis. Two separate PD controllers are used to calculate the two theoretic corrective servosphere speeds 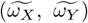 in addition to 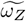 being kept at zero. These three values are transformed into the three rotor reference frames to obtain the signal that will generate the motion compensation.

### 3.2 Insects

Colonies of queen-right wood ants (*Formica rufa* L.) were collected from woodland (Broadstone Warren, East Sussex, UK) and housed in large tanks in the laboratory in the University of Sussex at 20–25 °C. Water, sucrose and dead crickets were provided *ad libitum* on the surface of the nest. To increase the ants’ foraging motivation, food was kept at a minimum during the experiments but ants had access to water all the time.

### 3.3 Testing innate behaviours

To evaluate any effects of the treadmill on the movement pattern and motivational state of ants, we conducted a series of experiments in simple environments. First, we surrounded the servosphere with a homogeneous white wall 10*cm* from the center of the sphere. We measured two main characteristics of the ants’ paths: the transition rates between active and inactive phases of movement; and the turning rate, represented by the angular speed. The statistics measured on the treadmill were compared with statistics measured from ants exploring a 120*cm* diameter circular arena surrounded by an uniform white wall. Secondly, we added a black vertical bar to the environment, to trigger directed behaviour towards the single conspicuous cue as previously observed in many insect species (Wallace, 1962; Voss, 1967; Wehner, 1972; Collett, 1988; Graham et al., 2003).

### 3.4 Testing associative learning

Twelve ants were tested (so-called pre-training group) on the treadmill to evaluate their innate preference for two different patterns made of red cardboard: a red circle; and two triangles merged together (figure 3C). Then, training was conducted in a T-maze connected to the ants’ nest-box (including individuals from the pre-training group and additional ants naive to these patterns). At the end of one arm of the T-maze we placed the circle pattern along with a feeder (50% water - 50% sugar). The arm in which it was placed was switched for every training session. Each session consists of 12h free exploration of the T-maze, after which the ants were collected and either tested or moved back into the nest-box. Ants had no access to food between each exploration session, to maintain their motivation. Post-training ants were tested on the treadmill using both patterns at once, in a competitive manner. The position of each pattern was kept constant during the trials on the treadmill (circle at —45°, two-triangles at +45°; figure 3C). The post-training group included pre-tested ants and ants that had never experienced the treadmill.

**Figure 3:**
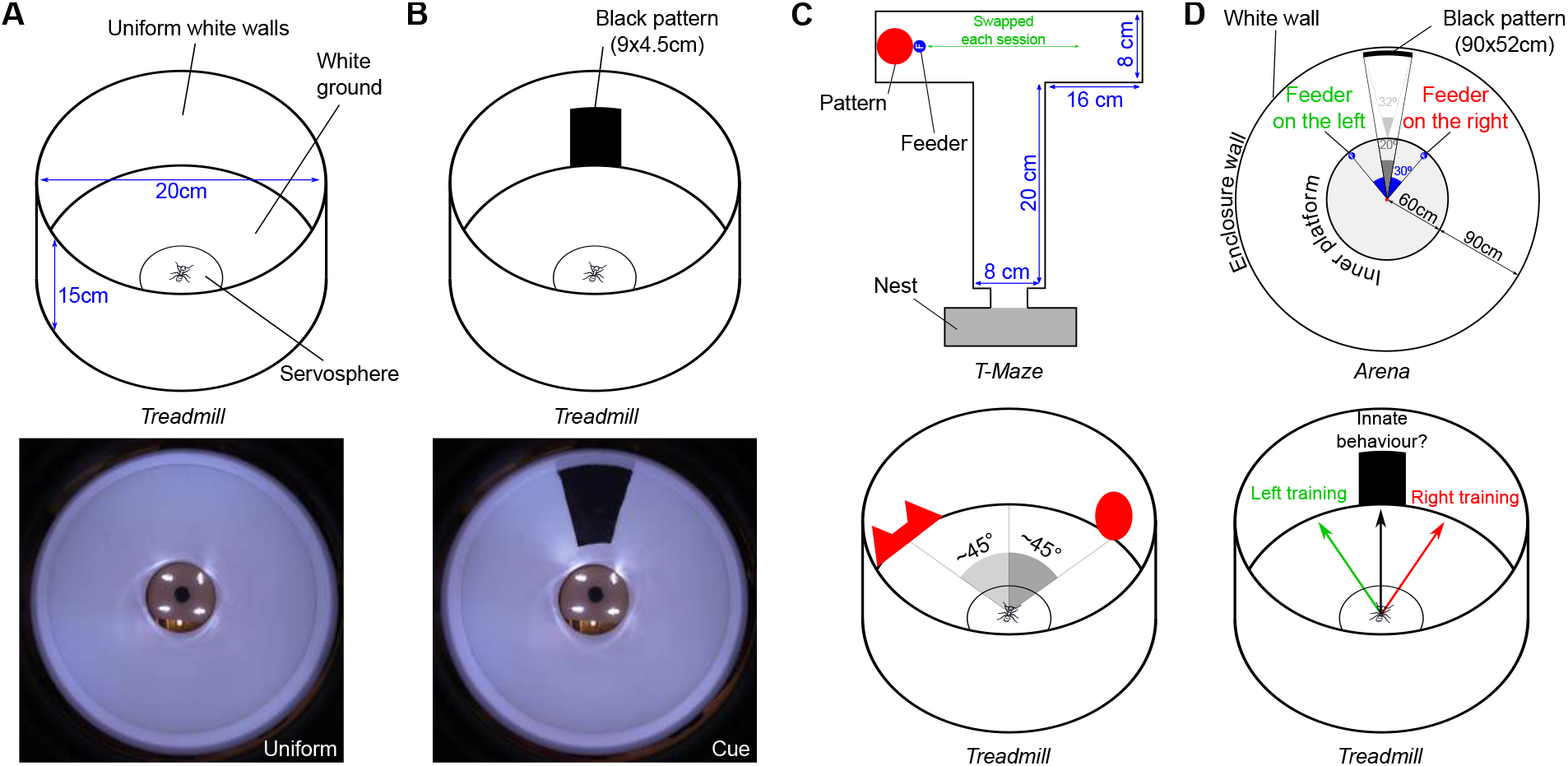
Schematic of the behavioural protocols. (A) Innate movement patterns were recorded on the treadmill in a circular arena of 20*cm* diameter and 15*cm* height. The inset picture at the bottom shows the visual environment from the ants’ point of view on the servosphere. (B) Innate response to a visual cue (a vertical black bar) was tested on the treadmill. The inset picture at the bottom shows the environment with the visual cue from the ants’ perspective. (C) Protocol used to test transfer of associative learning onto the treadmill. *Upper panel* Circle pattern was reinforced by systematically associating it with food reward in a simple T-maze. The pattern was placed in a different arm of the maze for each training session. *Lower panel* Preference for the pattern was tested in competition to assess the effect of the reinforcement. The patterns were always presented at the same location with respect to the center of the servosphere. (D) Protocol used to test the transfer of navigational memory onto the treadmill. *Upper panel* A subset of ants have been trained in an arena (150*cm* in radius) to find a feeder located 30° to the right or left of the outside of a black rectangle placed on the white circular wall of the arena. *Lower panel* Ants that reliably reached the feeder in the arena are tested on the treadmill for around 10 min.

We did not have information for individual ants about their degree of training in the T-maze. Nevertheless, each tested ant had been observed to leave the nest-box during at least one training session and therefore had potentially explored the T-maze.

### 3.5 Navigational task

A group of ants were selected as foragers based on their motivation to explore a cylinder placed around 20*cm* from a release point. These ants were individually marked and then trained in an arena to find a feeder displaced at a constant angle from a single, more distant, visual cue (figure 3D). The arena consisted of a 120*cm* diameter platform surrounded by a white cylinder of 300*cm* diameter and 180*cm* height. The visual cue was a black rectangle (52*cm* wide, 90*cm* high) placed on the outer wall. Throughout the training protocol, a feeder (67% water - 33% sugar) was placed at the outer edge of the central platform, offset by 30° to the right or left of the outside edge of the black rectangle.

Each ant experienced several training sessions, initially in groups of 5-6 ants, progressing to individual training. Ants were released in the center of the central platform at the beginning of each session and allowed to explore until they reached the feeder, where they could feed *ad libitum*. After they leave the feeder, ants had a small amount of time to further explore the surroundings before being returned to the nest where they can express trophallaxis with the rest of the colony. Ants showing foraging activity on the surface of the nest would be collected for a new training session. In total, including both collective and individual training, a training day included around ten sessions. After 2 to 3 days of training, the best trained ants, characterized by direct paths to the feeder on at least 3 sessions in a row, were selected to be tested on the treadmill.

On the treadmill, ants were tested with a vertical bar of 4.5*cm* width and 8*cm* height placed 10*cm* away from the center of the servosphere (25°x38.5°), which matches approximately the angular size of the cue experienced by the ants near the start of their trip from the center of the arena (20°x31°) to the edge of the platform (60°x71.5°). The visual cues used in both contexts have the same 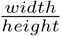 ratio (Treadmill: 0.5625; Arena training: 0.5778). Each ant was tested once for 10 minutes after which it was placed in a tank providing food and then moved back to the nest. We then made a further test of these ants in the familiar arena, to check for any change in their motivation to forage or extinction of their memory due to the testing on the treadmill.

## 4 Results

### 4.1 Innate behaviours

One of our aims in developing an untethered treadmill for ants was to avoid artefacts caused by tethering and, hence, to be able to observe robust naturalistic behaviour in virtual reality. However, the potential vibration produced by controlling the 3-DOF servosphere could also cause some disturbance that modifies the ants’ behaviour. In order to measure the consistency of ant movements we conducted a set of experiments to quantify innate behaviour on the treadmill in comparison to conventional arena experiments.

First, we placed ants in an uniform environment to measure motion statistics on the compensator and in an arena (Figure 4). Wood ants showed surprisingly robust exploratory behaviour during long periods of time on the treadmill, including many 15 to 30 minute bouts without significant drops in activity, and one ant running for 150 minutes. This suggests an advantage of the untethered setup compared to some tethered conditions where the insects’ motivation can drop quickly (personal communication). Ants surrounded by a uniform white environment (figure 3B) explored the entire extent of the virtual arena although they expressed a slight preference towards one direction (Omnibus test: *z* = 5, *p* = 0.020; Figure 4B). To look in more detail at the movement of ants on the treadmill we selected 2 main parameters of interest. The first parameter we measured represents the tendency of wood ants to move in a stop & go manner. We measured the distribution of the lengths of each of the walking (active) and pausing (inactive) bouts. We observed a fairly similar distribution between arena and treadmill experiments in term of active and inactive bout length (figure 4C & figure A.2BC), showing that this movement pattern is not affected by our setup. The second parameter we looked at is the distribution of angular speeds, as a turning-rate proxy. On both the treadmill and in an arena ants displayed the same range of angular speed in-between —200 and 200°.s^−1^ (Figure 4D & figure A.2A), although we could observe an increase in the peak at 0°.s^−1^ on the treadmill, i.e. a higher tendency to walk straight. This may be explained by the lack of any physical constraints on direction of progress on the treadmill, whereas ants in the arena will be forced to turn when reaching the edge.

**Figure 4:**
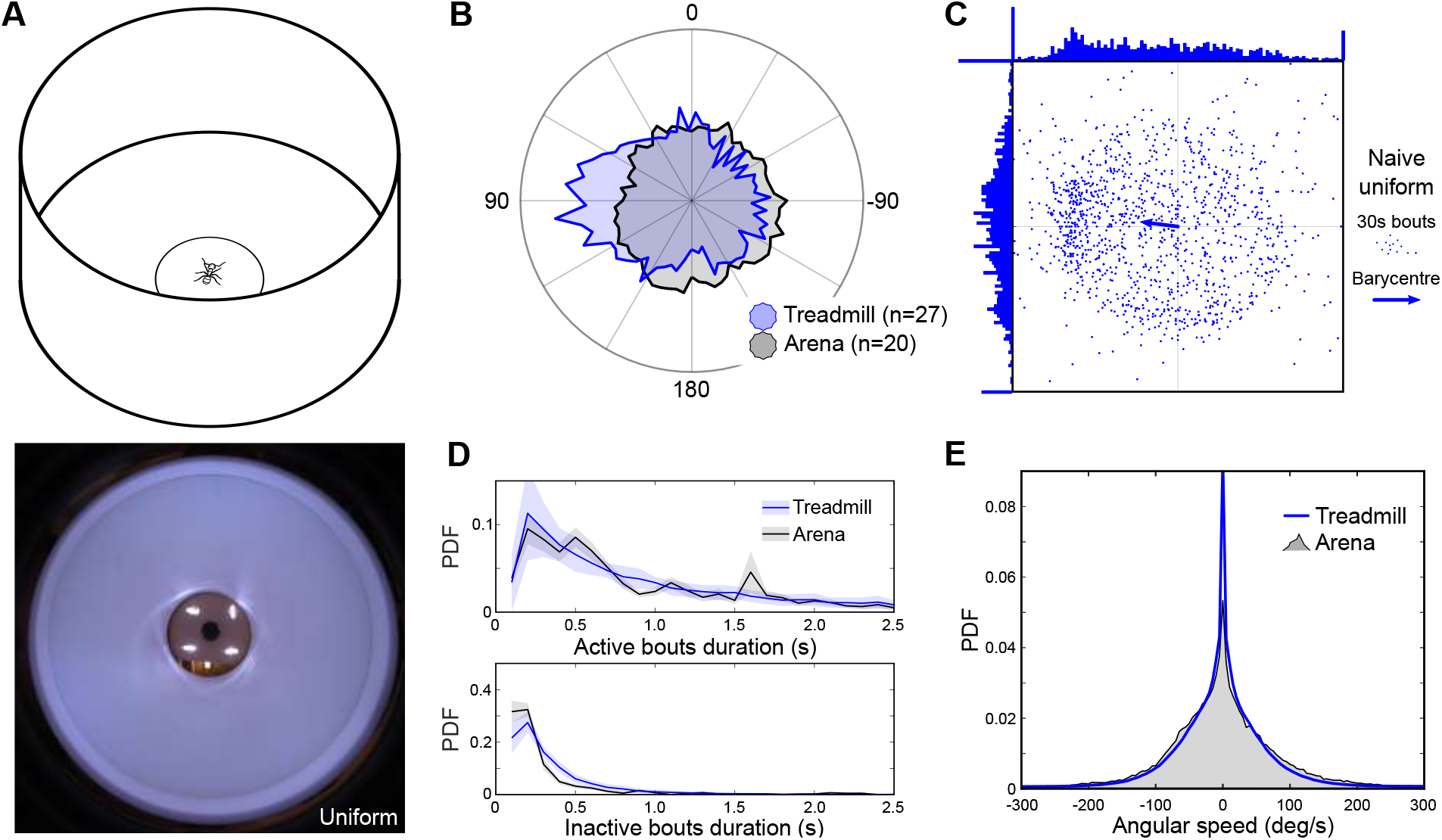
Walking dynamics in an homogeneous white environment on the treadmill correspond to the arena condition. (A) Conservation of ants’ innate motor patterns has been tested on the treadmill in a circular arena of 20*cm* diameter and 15*cm* height. Lower panel shows the visual environment from the ants’ point of view on the servosphere. (B) Probability density function of the heading measured on the treadmill (blue patch) and in an arena (black patch) both in an uniformly white environment. The distribution of headings onto the treadmill shows a statistically significant bias from uniformity [Omnibus test: Arena - z = 4, p = 0.11; Treadmill - z = 5, p = 0.02]. (C) Mean resultant vector extracted from the experiment in an uniformly white environment on the treadmill. Each point represents the mean resultant vector of movement for a 30s bout (i.e. vectorial sum of all the unitary movements during 30s). The arrow indicates the overall resultant vector, showing there is a small bias, mostly along the Y-axis, in the uniform arena (Omnibus test: z = 5, p = 0.020). (D) Distribution of active and inactive bout length during the experiments on the treadmill (blue line) and in the arena (black line) in an uniformly white environment. (E) Distribution of angular speed measured during the experiments on the treadmill (blue line) and in the arena (black patch) in an uniformly white environment.

In addition to checking for the resilience of the innate movement pattern, we looked at whether innate directed behaviour towards a single visual cue Wallace (1962); Voss (1967); Wehner (1972); Collett (1988); Graham et al. (2003) is maintained on the treadmill. Ants were presented with a vertical black bar with an angular size of approximately 20° from the center of the treadmill in a totally white environment (figure 3B). We clearly observed an oriented behaviour towards the cue for each individual when compared with the uniform environment (figure 5B). We also observed that the movement pattern of the ants, both in term of active-inactive bouts and turning rate, was not drastically affected by the presence of the cue (figure 5CD & figure A.2).

**Figure 5:**
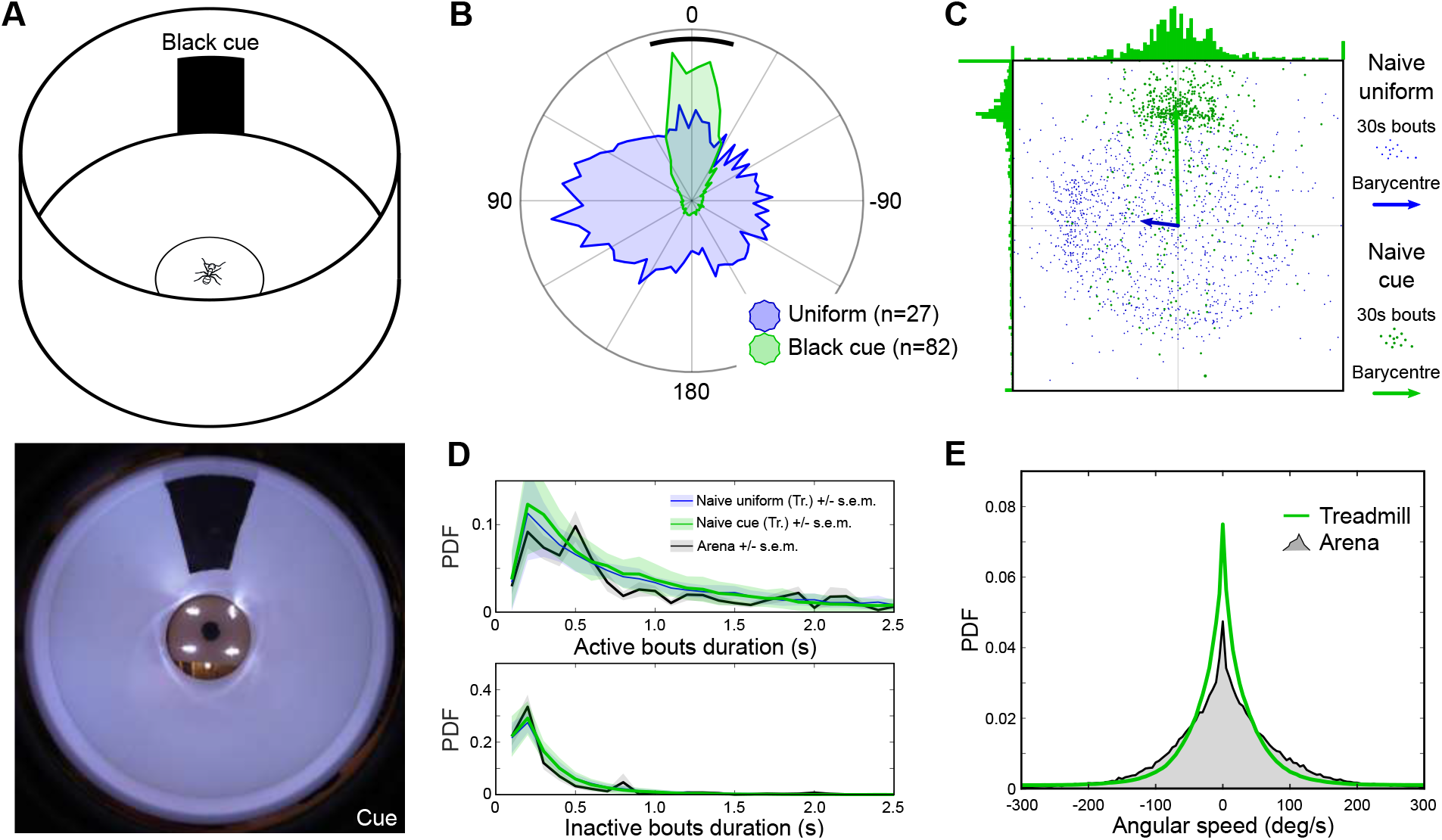
Innate attraction toward a single conspicuous cue on the treadmill correspond to arena condition. (A) Innate response to a visual cue (a vertical black bar) has been tested on the treadmill. The inset picture at the bottom shows the visual environment with the added cue from the ants’ point of view on the servosphere. (B) Probability density function of the heading measured on the treadmill in an uniformly white environment (blue patch) or in the presence of a visual cue (green patch). (C) Mean resultant vector extracted from the experiment in an uniformly white environment (blue points) or in presence of a cue (green points) on the treadmill. Each point represent a 30s bout mean resultant vectors of movement. The arrows indicate the overall resultant vectors for each group. The cue induce a clear and important bias over the X-axis and overpass totally the bias observed in the uniform white environment. (D) Distribution of active and inactive bout length during experiments on the treadmill with a single cue (green line) or in an uniformly environment (blue line) and in the arena with a corresponding cue (black line). (E) Distribution of angular speed in experiments with a single cue on the treadmill (green line) or in the arena (black patch).

### Transfer of learned behaviour

Our innate behaviour tests showed that ants maintained a relatively normal motor pattern on the servosphere despite any vibrations and the discrepancy in visual feedback implied by walking on the spot, but it did not allow us to check for the robustness of the motivation to express more complex behaviour such as foraging. To test this, we conducted 2 different experiments.

We first used the ability of insects to associate a visual cue/pattern with reward (Guerrieri and d’Ettorre, 2010; Schwarz and Cheng, 2010, 2011; Giurfa, 2012) (Figure 3A). Before any training, ants expressed a slight attraction to the two-triangles pattern (Figure 6A equivalent t-test: *μ_pre_* = 15.04;, 95% confidence interval [2.03 28.05]). We therefore decided to reinforce the other cue during the training in a T-maze in order to better identify its effect on the choice of the ants on the treadmill afterwards. Indeed, after training, ants showed a clear shift toward the circle cue (Watson-Williams test: *F* = 9.56; *p* = 0.0038; *μ_post_* = −21.06, 95% confidence interval [−37.42 −4.69]), suggesting a transfer of memory from the T-maze context, as well as the preservation of foraging motivation on the treadmill.

**Figure 6:**
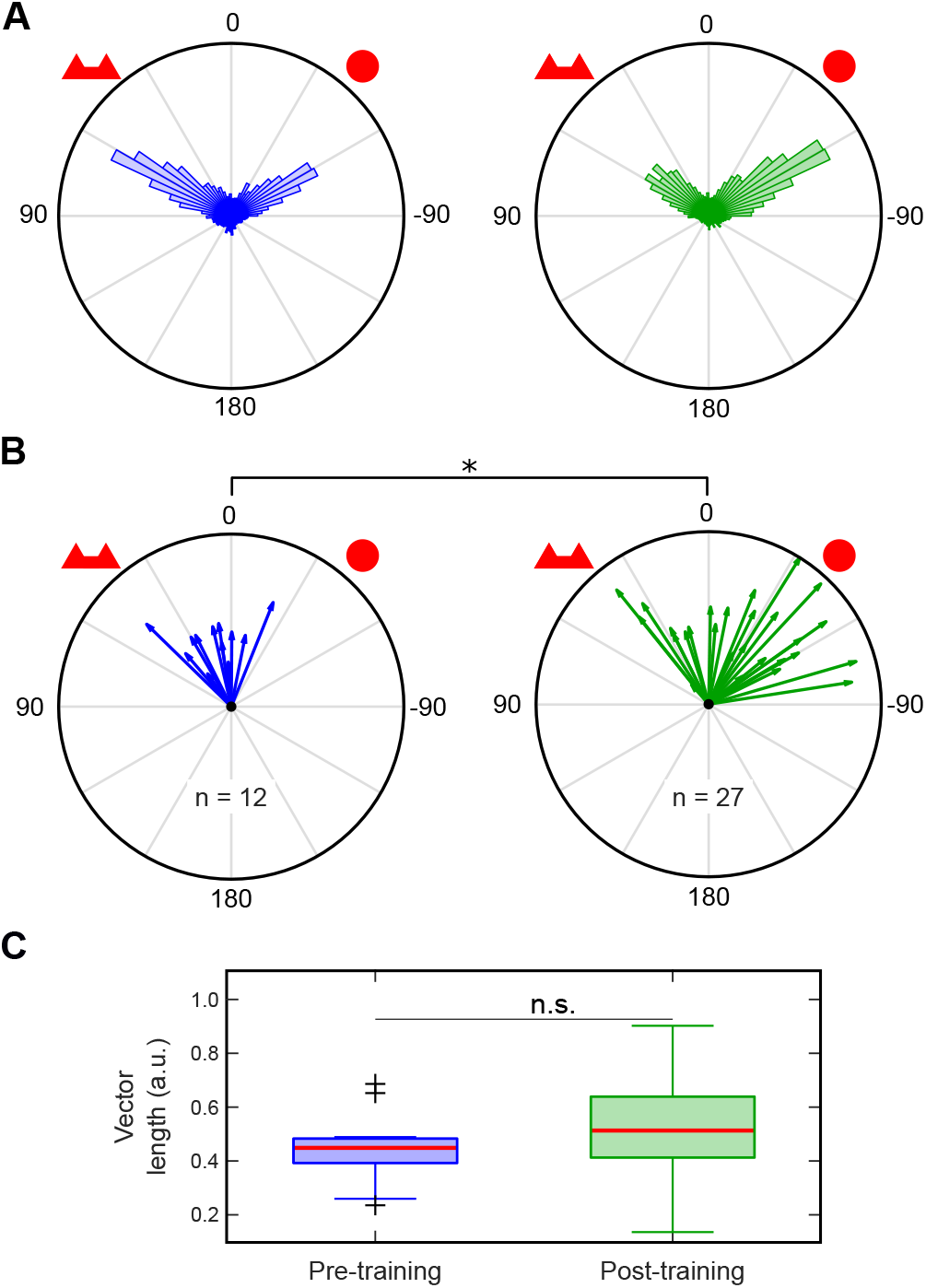
Associative visual memory acquired during a T-maze conditioning is transferred onto the treadmill. (A) Distribution of pooled heading data in both pre-tested (blue) and post-tested (green) ants. (B) Arrows indicate the orientation of the resultant mean vector of each experiment in pre-tested (blue) and post-tested (green) ants. Distribution of the vectors is significantly different between pre- and post-tested ants (Watson-Williams two-sample test: *F* = 9.56, *p* = 0.0038). (C) Boxplots of the length of the resultant mean vector for each trial of both groups (no statistical significant difference between the groups, Kruskal-Wallis test: *Chi*^2^ = 1.34; *p* = 0.2476). *: *p* < 0.05; n.s.: not significant (*p* > 0.05).

Going beyond simple association of a cue with reward, ants are able to learn the location of a feeder relative to distant visual cues Durier et al. (2003). We assessed the ability of ants to reproduce such a navigational task on the treadmill by training them to reach a feeder displaced from a vertical bar in a circular arena of 120*cm* diameter (figure 3B). To focus only on the transfer of the memory and not on the intrinsic ability of ants to perform the task, we first selected the ants that showed the best performance during training in the arena, by selecting those ants that showed fairly direct paths to the feeder for several training sessions in a row (figure 7A). Trained ants took less than 40s to reach the feeder at the edge of the central platform (≈ 60cm for a direct path). Note that the distribution of the feeder position in the visual field of the ants does not appear to reflect a strategy by which the visual cue is kept in a constant angular position, but rather that it oscillates between 0° and ±60°, depending on the training group (figure 7A). This is confirmed by a more detailed analysis of the dynamics of the ants’ headings, which oscillate around the feeder location (figure 8A) and often use fixation of the cue as a “limit” of the oscillation.

**Figure 7:**
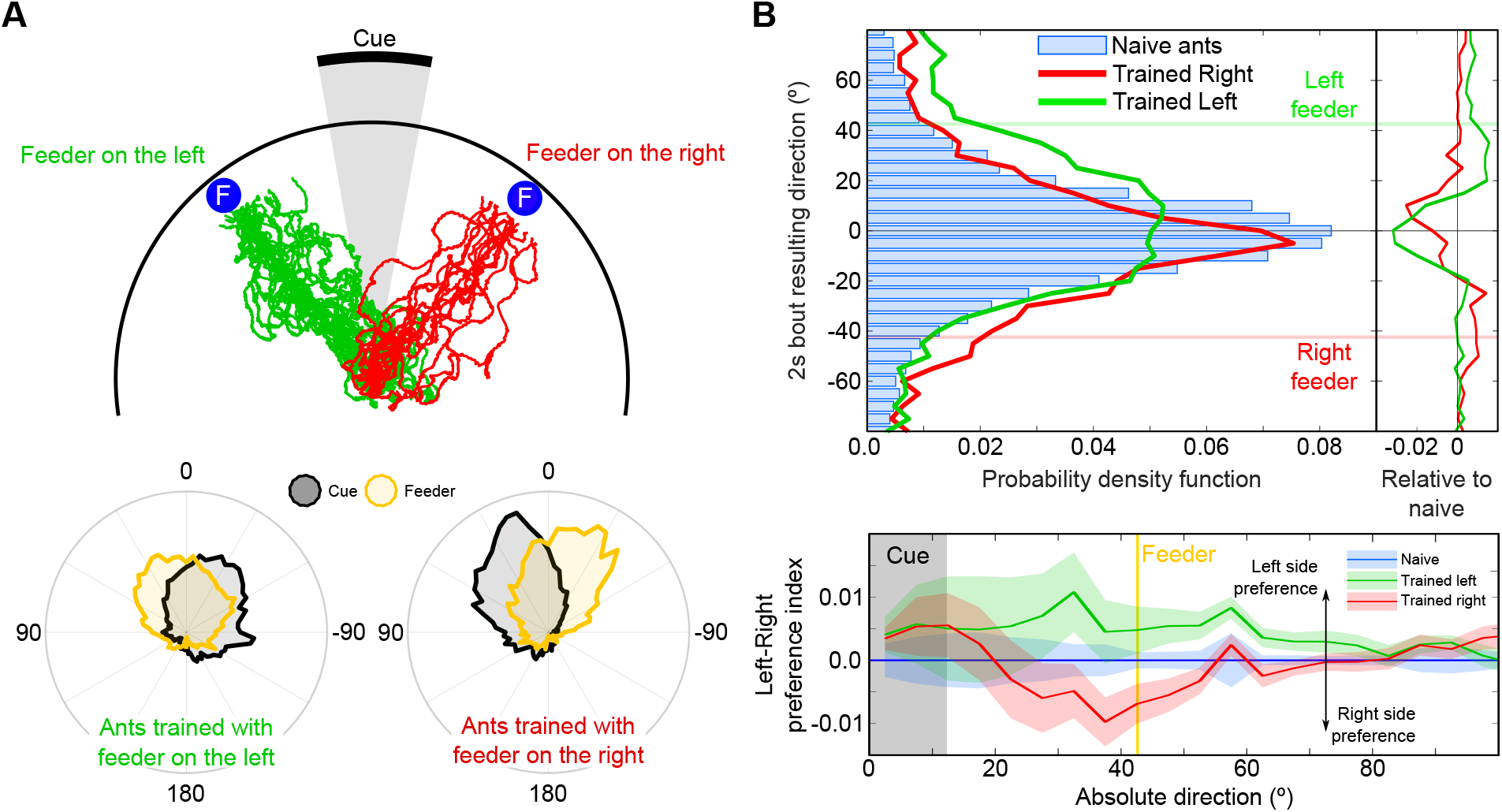
Navigational memory of a feeder location acquired during arena conditioning is transferred onto the treadmill. (A) *Upper panel* Paths of the selected ants during the last arena (120 cm in diameter) training before they were tested on the treadmill. *Lower panel* Distribution of both the visual cue and the feeder in the field of view of the ants trained with the feeder on the right or on the left of the cue during their last training session. (B) *Upper panel* Distribution of the orientation of the mean resultant vectors calculated for every 2*s* bout during the first five minutes of each recording for naive ants (blue bars) or for trained ants with the feeder on the left (green line) or on the right (red line) of the cue. The right plot shows the same distributions for the two trained groups relative to the naive group distribution (Naive distribution subtracted). *Lower panel* Left-Right mean preference index for groups of ants trained with the feeder on the left (green) or on the right (red) of the cue. For each trial, the index is calculated by comparing the probability function the difference between opposite orientations for each absolute angle relative to the cue *α*: *index*(|*α*|) = *p*(*α*) − *p*(−*α*). A positive value indicates then a preference for the left side of the cue and a negative value a preference for the right of the cue. The index calculated for the naive ants (blue shaded area) has been subtracted from each group of trained ants. Shaded area indicate the standard error of the mean.

When tested on the treadmill, trained ants oriented themselves rather more toward the cue than the feeder (figure 7B), although the training induced a significant effect in the direction of the virtual position of the feeder (Williams-Watson test: *F* = 6.65; *p* = 0.0013). This trend was small but when compared with the naive distribution of orientations, a clear bump appeared around the location of the right or left feeder for each corresponding trained group (figure 7B). Furthermore, by calculating a left-right preference index (figure 7B) in regard to the cue location we observed that the preference for the virtual feeder location was clearly determined by the training condition (either −42.5° or 42.5° from the centre of the cue). It is also noteworthy that in each group the behaviour of the ants in the neighborhood of the cue clearly expressed a preference for the right edge of the cue [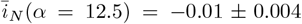 *s.e.m*. (Wilcoxon test: *z* = −2.7714; *p* = 0.0056); 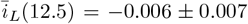; 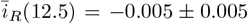; with 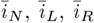 the mean preference index for respectively the naive group, the group trained with the feeder on the left, and with the feeder on the right].

Figure 8 shows a comparison of the heading dynamics in the arena at the end of the training sessions (figure 8A) and on the treadmill (figure 8B). On the treadmill (figure 8B), the oscillation of the heading was nicely conserved and we can observe how the headings tend to get centered either on the cue or in-between the cue and the feeder orientation. Nevertheless, the ants had an initial period of adaptation to the treadmill, of around 20s in the 2 examples presented (figure 8B1 & B), during which the heading was clearly more erratic, probably indicating a necessary delay for the ants to settle to this unfamiliar environment. This adaptation period was of the same range of time for each ant as shown in figure 8. Except during this period of potential adaptation to the context, ants seem to express comparable strategies in the exploration of their environment (figure 8B3). Oscillations are fairly similar in terms of amplitude (signals have been purposely centered but not normalized) and only slightly different in term of frequency.

**Figure 8:**
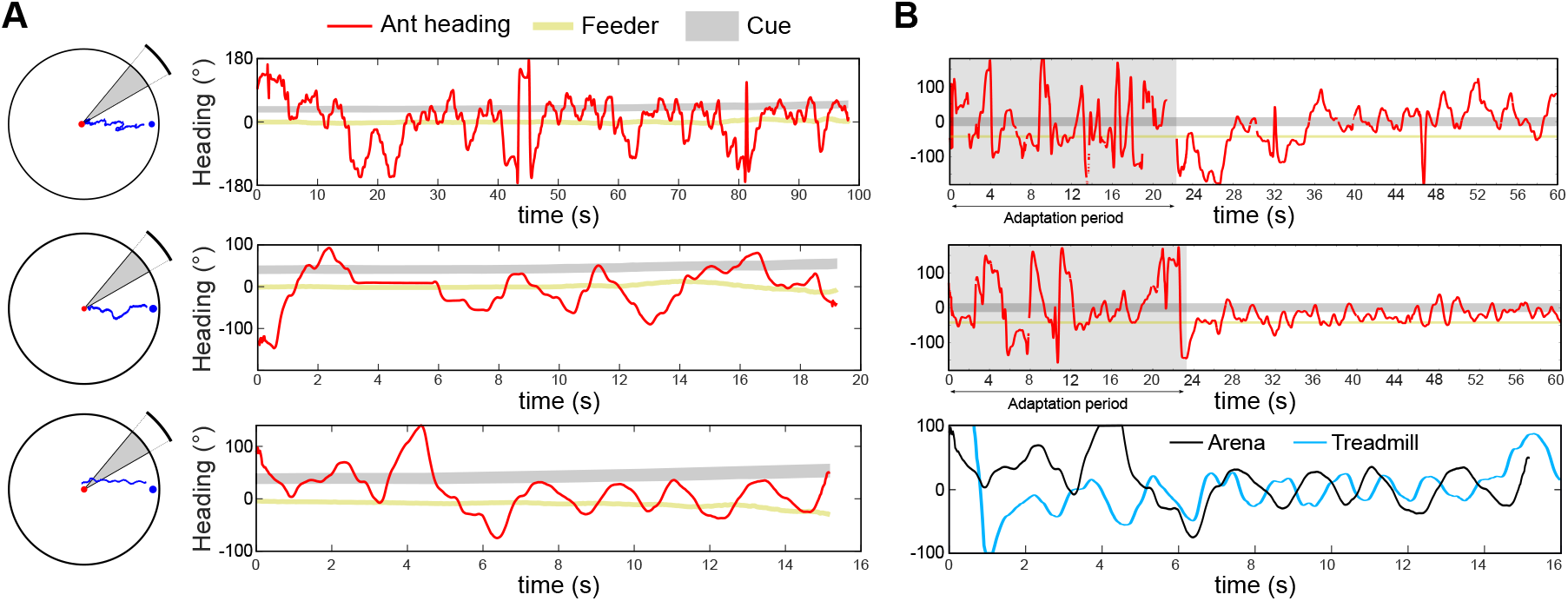
Oscillatory heading behaviour in training ants is transferred from the arena to the treadmill. (A) Examples of heading dynamics during the training in the arena (red trace). The black band shows the evolution of the orientation of the cue in regard to the ant’s position along the path and the gold line the orientation of the feeder. On the left are shown the XY positions of the corresponding path. (B) The 2 first panels show examples of heading dynamics during the first 60 seconds on the treadmill of the corresponding ants. The last panel shows both heading dynamics in the arena (black line) and on the treadmill (blue line) during 16s. The treadmill path has been selected starting just after the ≈ 20*s* of adaptation observed generally (see panels above). Both signals have been centered arbitrarily to compare oscillation dynamics, but not normalized (i.e. the amplitude is conserved).

Finally, to complete our experiments, and because we are aiming at a device that allows interaction between classic arena/maze/field and VR experiments, we tested whether experiencing the treadmill would disturb memory or behaviour during retrieval experiments. At the very least, the total absence of reward during the trials on the treadmill could induce a reduction of the memory, or even reversal if this is experienced as punishing. To test this, we placed ants that had been tested on the treadmill back in the arena for an additional ‘‘training” session. Figure 9A shows the ants’ path when experiencing again the cue in the arena. The directions taken to reach the feeder were clearly more noisy compared to the one at the end of the training (figure 7A & C; Bartlett’s test: *T* = 13.893, *p* = 0.0002) but still centered towards the feeder location (figure 7C; Kruskal-Wallis test: *Chi*^2^ = 0.05, *p* = 0.82), rather than attracted by the cue as we would expect from a return to naive state. Furthermore, the pattern of fixation of both the cue and the feeder location did not change drastically after unrewarded experiments on the treadmill (figure 9B) and the time taken to reach the edge of the platform did not seem to be much delayed (figure 9D; Wilcoxon test: *p* = 0.81), suggesting straight path toward the edge of the platform and a maintained foraging motivation. These retrieval experiments were only conducted for ants from the group trained with the feeder on the right of the cue. Nevertheless, in the group trained with the feeder on the left, ants selected first to be tested on the treadmill were allowed to rejoin the training afterward and therefore could be selected several times. The observation of multiple selection (2 or 3 times) for several ants did not appear to alter the behaviour, supporting the paradigm of back and forth transfer between real arena experiments and virtual reality treadmill experiments. Overall, it appears that the experience of very long (10*min*) unrewarded trials on the treadmill in comparison to ≈ 40*s* rewarded trials in the arena did not significantly extinguish the memory or the motivation of the ants.

**Figure 9:**
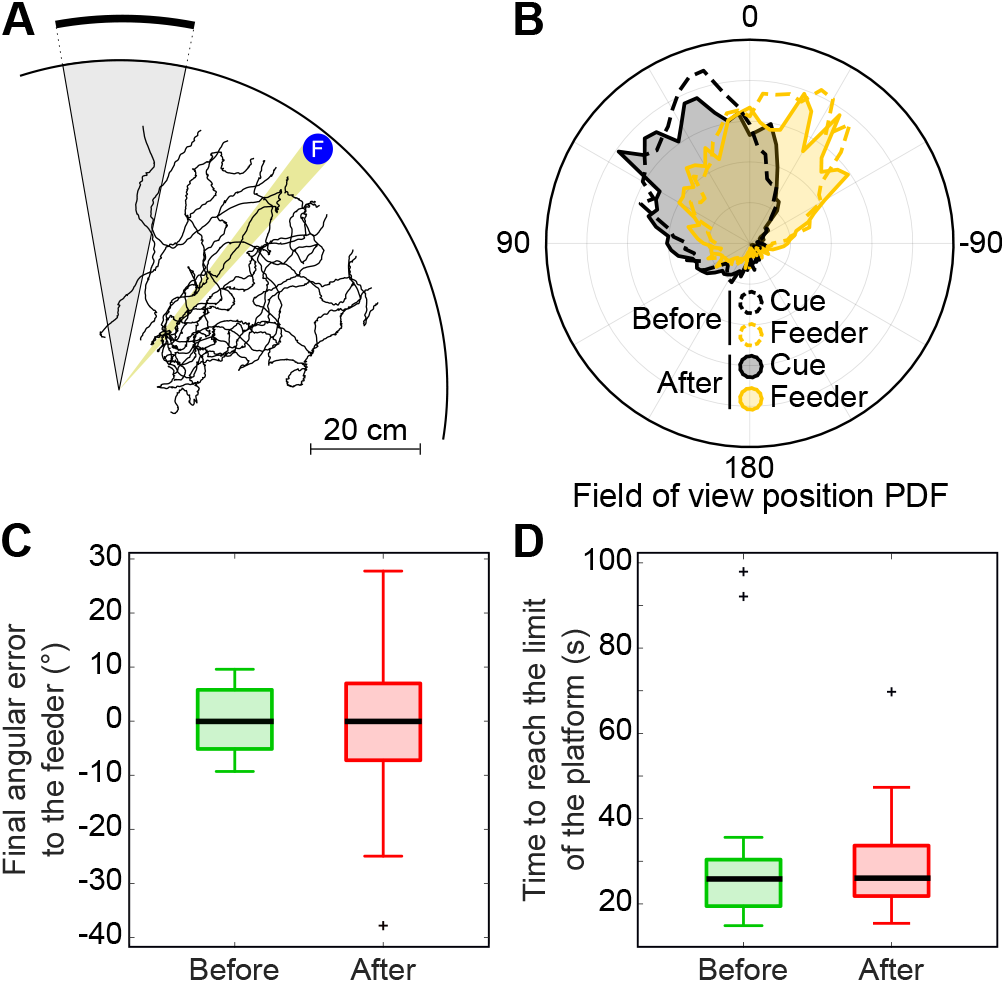
Feeder location memory is preserved after experiencing long unrewarded experiment on the treadmill. (A) Paths observed in the arena directly after the experiments conducted on the treadmill. (B) Distribution of the orientation, relative to the ant’s orientation, for the cue (black) and the feeder (yellow) before (dotted lines) and after (solid lines) the test on the treadmill. (C) Final angular error of the ant relative to the feeder calculated during the last learning session before (green) or the first learning session after (red) the test on the treadmill. (D) Time to reach the edge of the inner platform (estimated at 50*cm* from the center of the arena) calculated during the last learning session before (green) or the first learning session after (red) the test on the treadmill.

## Discussion

Our aim was to develop a means to study insect visual navigation in well-controlled but naturalistic walking conditions. To this end, we developed a treadmill (a 3 degrees of freedom (DOF) servosphere) for wood ants (*Formica rufa*) that compensated for their movement such that they remained in one location relative to their surroundings whilst freely walking untethered. Such treadmills have been employed previously in studies of insects with slow dynamics of movement (earlier work reviewed in (Hedwig, 2017), more recent examples include (Sakuma, 2002; Shigaki et al., 2016)) but not for faster insects such as ants. We showed that using visual feedback control the ant can be kept reliably within 5 mm of the apex of the sphere. We also showed that ants on the treadmill maintained similar walking dynamics to freely moving ants walking in an arena (figure 4 & 5), and could transfer learnt visual associations between the two contexts (figure 6 & 7).

### Advantages and disadvantages of the motion compensator

Many studies have used trackballs to investigate in detail how animals move in relation to a sensory stimulus such as an olfactory, auditory or visual cue (Schultheiss et al., 2017). These trackballs typically involve a sphere suspended on a cushion of air that is propelled by a tethered animal atop it (Van Swinderen, 2012; Moore et al., 2014; Paulk et al., 2014; Dahmen et al., 2017). This can set strong constraints on the sphere size, weight and shape, typically requiring bespoke manufacture of the system (Dahmen et al., 2017). A practical advantage of the motion compensator is that it can be built from commercially available components, and larger spheres (with lower curvature) can be used because the sphere is moved by motors not the insect.

An additional advantage is that the motion compensator does not require a tether to be attached to the insect. This reduces or removes: (i) the effects of handling, (ii) the introduction of undesirable variability dependent on the expertise of the experimenter, and (iii) the potential change in motivation and walking dynamics caused by the attachment of a tether. It is known that handling of insects induces stress and the release of neuromodulators which can alter behaviour long after the cause of the stress ceased (Davenport and Evans, 1984; Harris and Woodring, 1992; Roeder, 1999). Recent studies also shown how aversive experiences can quickly affect the behaviour of ants (Wystrach et al, 2020). The diminished need for manipulation of the animal also facilitates the transition between naturalistic field work and fully controlled laboratory approach. Complex navigation experiments with social insects require the ability to return ants to their colony to feed their conspecifics in order to maintain their foraging motivation. Our apparatus allows a combination of field/arena/treadmill experiments to be used at different stages of a training program, for example. In addition, behaviour persisted over long durations, over 2 hours in some cases, and thus could support experiments more consistent with the long distances central foraging insects can cover.

Tethering insect on a trackball is that tethering can directly affect both the motivation and the motion of the animal (Spirito and Mushrush, 1979; Taylor et al., 2015). Our treadmill obviates the need for the physical constraints of a tether allowing more naturalistic movement of the wood ants, as observed on a similar motion compensator for pillbug (Nagaya et at., 2017). Although some aspects of the motion compensation (such as vibration or altered motion feedback) could have altered the ants’ behaviour, our results suggest that walking dynamics were not drastically impacted by the motion compensator. Comparison between ants on the treadmill and in an arena show no change in several aspects of walking such as the distribution of walking bout durations and the angular speed in a uniform environment and with a salient cue. In our experiments, behaviour persisted over long durations even though walking on the spot is an open-loop situation and therefore should elicit some discrepancy in feedback, which could have led to a decrease in motivation. It still needs to be shown that additionally correcting for the ants’ angular velocity to keep them in a fixed orientation, as can be done on the compensator (Shigaki et al., 2016), would not cause a similar decrease in behavioural motivation as seen in classic trackball experiments that prevent free rotation. (Buatois et al., 2017)

The obvious disadvantage of not tethering the insect is that it prevents simultaneous intracellular neural recordings or imaging of brain activity, which is possible from head-fixed insects on trackballs. Nevertheless, extracellular recordings and optogenetic stimulation should remain viable and easier to achieve than in an arena due to the limited movement distance.

### Transfer of learnt behaviour on the motion compensator

Showing that the walking dynamics are not radically altered by the active motion compensation provided by our treadmill does not establish its suitability for navigation experiments. In particular, these require that the ant on the treadmill responds to presented stimuli in a manner consistent with its previous training experience off the treadmill. Therefore we developed two different experiments to inspect the transfer of learnt tasks on the treadmill, an associative learning task and a proper navigation task.

Ants trained in a T-maze to associate a visual pattern to a food reward showed clear preference for that pattern when tested on the treadmill (figure 6). Transfer of learning in different contexts is a known ability in insects. It has particularly been shown in VR setups using associative learning paradigms (Buatois et al. (2017); Rusch et al. (2017) reviewed in Buatois (2018)). Here, it suggests that the recognition of certain visual cues can occur independently of the panoramic surrounding. Furthermore, our treadmill, by giving the ants the ability to freely scan the environment, allows a more detailed observation of active learning or recognition strategies (see in figure 8). Recording orientation behaviour in the continuous manner offered by the treadmill (rather than at a single choice point) allows observation of persistence or suppression mechanisms in the memory process. In addition, long duration observation provides a stronger statistical power to evaluate dichotomous and sequential choice in ants (Chameron et al., 1998; Schatz et al., 1999; Beugnon and Macquart, 2016).

Ants trained in an arena to approach a feeder displaced to one side of a visual cue (a vertical black bar) showed a tendency to orient their exploration toward the virtual position of the feeder (figure 7B) when tested on the treadmill, again showing the ability of ants to use certain cues independently of the complete visual scenery. It also support the preservation of ants foraging motivation on the treadmill, crucial point in respect to experiments on navigation behaviour. However, the shift in direction displayed on the treadmill was relatively weak. There may be several reasons for this. Firstly, the absence of any change in the size of the cue, and consequent discrepancy in the expected optic flow, could affect the behaviour (Brembs, 2009). Secondly, we observed that in the arena, orientation to the feeder did not occur immediately after release in the centre (figure 8A), suggesting that the cue could initially directly serve as a target (Graham et al., 2003). Therefore, our use on the treadmill of a cue with an angular size close to the cue observed from the centre of the arena may not have fully elicited the trained deviation towards the feeder. Thirdly, the strategy used by ants to reach the feeder in the arena is not to keep a constant heading towards the feeder location, but to oscillate around its orientation (Collett et al., 2014), and moreover to use the cue as a limit of the oscillations (figure 8A). It is noteworthy that the exploratory oscillations are preserved on the treadmill. Although we only looked in detail at a few trials (figure 8), the systematic analysis of the ants’ paths in a fully controlled closed-loop visual environment could be crucial to understand the active strategy used by ants to inspect, memorize and recognise their paths.

A surprising result we observed during this experiment is the existence of a seemingly innate preference for the right edge of the cue (figure 7B). This right-edge preference occurs for ants trained with the feeder either on the right or on the left, suggesting a robust bias. Lateralisation in the insects’ brain has received an increasing attention in the last decade, particularly from an adaptive perspective (Niven and Frasnelli, 2018; Frasnelli, 2013), and in wood ants it has been observed in behaviour such as trophillaxis (Frasnelli et al., 2012) or memory encoding (Rogers and Vallortigara, 2008).

### Potential applications of the treadmill

The existence of transfer of associative memory from classic T-maze to the motion compensator not only offers perspective to study memory processes but also provide an ideal benchmark to assess questions on upstream visual processes. The early processes at work in the visual pathways of ants have received little attention to date (Vowles, 1965; Cammaerts, 2008, 2013; Schwarz et at., 2011). Treadmills offer a tool for reverse engineering the visual abilities of ants in a more automated and large-scale approach than arena or maze experiments (Nagaya et at., 2017; Shigaki et al., 2016). Understanding feature extraction processes would be particularly crucial to incorporate in models that have been proposed to sustain visual memory of routes in ants for example (Ardin et al., 2016).

An obvious requirement to facilitate such work is the addition of a naturalistic and closed-loop virtual environment to the treadmill set-up. VR systems have already been shown to be a realistic solution to study insects vision in tethered (Dickinson, 2001; De Bivort and Van Swinderen, 2016) or in arena conditions (Fry et al., 2009; Straw et al., 2010). The combination of a 3D virtual environment and the preservation of the naturalistic walking behaviour on a motion compensator should allow fully monitored training and experiments, accelerating our understanding of how insect navigation is controlled by the brain.

## Acknowledgements

We are very thankful to Douglas Howie for the design of the treadmill mechanical parts and to Jan Stankiewicz for the fruitful discussions concerning the control and the electronics.

## Contributions

R.G. built the experimental setup (software and hardware). R.G., C.B., J.E.N., P.G., B.W. designed the experiments. R.G. and C.B. performed the experiments. R.G. analysed the data. R.G. wrote the initial version of the manuscript. R.G., C.B., J.E.N., P.G., B.W. reviewed and edited the manuscript.

## Data availability

Data are available at https://drive.google.com/open?id=1-du3-X8DgyeoLosv0vB6eETtO9SGvTbU

## Appendix

## 1 Hardware & software

### 1.1 Tracking system

The real-time communication with the camera has been ensured using python, a special wrapper pypylon (available on `PyPylonGithub` https://github.com/mabl/PyPylon) and the pylon camera software suite 5.0.12. The frame rate of the camera is maintained constant by processing the image extraction on a separated thread, which avoid any conflict with the main loop of the python code.

To improve tracking efficiency and maximize the rate of the overall feedback loop, the optics of the camera were slightly unfocused (figure 1B), thus acting as a physical filter instead of including it in the software. This is also helping by decreasing the background noise which can be critical for the functioning rate of the system at the contours detection stage. This background noise is additionally decreased by setting the binarization threshold to avoid any detection of the contrasts of the ball itself or of the potential shadows created by the lighting. All the features extraction are realized using openCV python package (pyopenCV).

In order to measure the precision of the system to keep the ants on the spot (figure 1), the error measurements has been converted from pixels 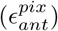 to millimeters 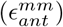 using the theoretical geometry of the setup [Distance of the camera: *D_cam_* = 150*mm*] and the characteristics of the camera [Pixels size: *S_pix_* = 4, 8.10^−3^*mm*; Focal length: *F_cam_* = 12*mm*]:

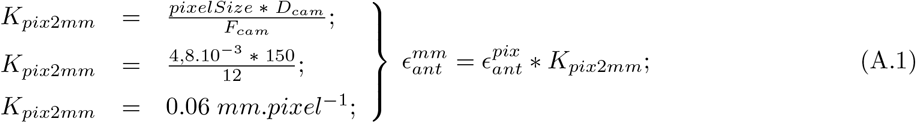

The calculate value of *K_pix2mm_* has been validated using a checkerboard and a custom calibration algorithm. The blurred image obtain with the unfocused lens did not allows us to use the built-in calibration toolbox from Matlab© for example. This validation showed slightly lower *K_pix2mm_* and therefore the geometry-based has been kept.

The subsequent control of the rotors is monitored by an Arduino© developmental board, using the HalfStepper library, smoothing the rotors motion, and ULN-2064 bridges. All the connections necessary to ensure the proper control of the rotors are presented in figure A.1.

**Figure A.1:**
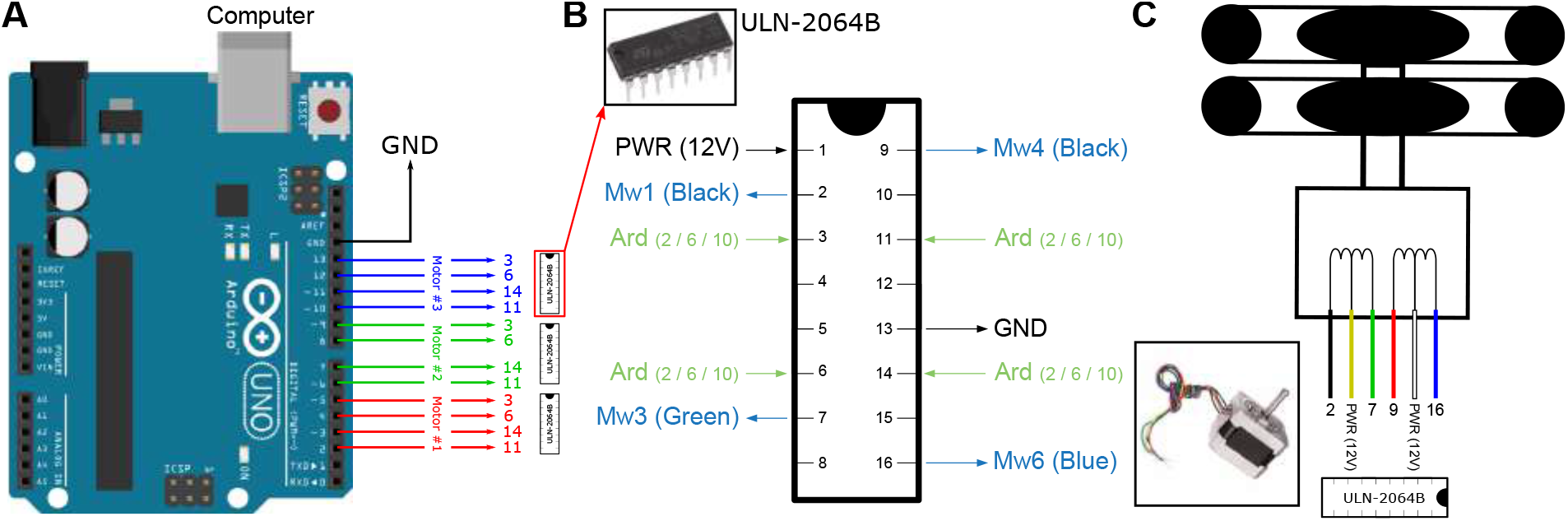
Connections from Arduino Uno to stepper motors. (A) Connection from the Arduino Uno. Each motor is connected via ULN-2064B to 4 digital pins: 1-4 for motor 1, 5-8 for motor 2 and 9-12 for motor 3. The ground as to be connected to the ground of the power supply used for the motor (12V). (B) Connection of the ULN-2064B with Arduino (green), to the motors (blue) and to the 12V power supply (black). Each motor is associated with a single ULN-2064B. (C) Stepper motors SY42STH38-0406A connections with its own ULN-2064B bridge. Yellow and white wires are connected directly with the 12V power supply. In case of different steppers used, you can check with the datasheet to adapt the connection based on their own wiring diagram.

## 2 Motion parameters comparison

To compare the distribution of the different parameters of the motion in-between the different condition (arena *vs* treadmill), we used the square root of the Jensen-Shannon divergence, also known as Jensen-Shannon distance. The Jensen-Shannon divergence [JSD(P—Q] between two probability density function (P and Q) is a symmetric and smoothed adaptation of the Kullback-Leibler divergence [D(P—Q)], defined by:

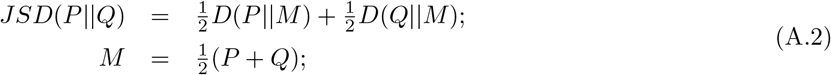

The square route of the JSD is in the interval [0; 1], nevertheless, as it is impossible to conclude statistically to the similarity between two different distribution we rather conducted the comparison of intra- and intergroup comparison to evaluate the evolution of this distance for the different parameters once the ants were placed on the treadmill.

**Figure A.2:**
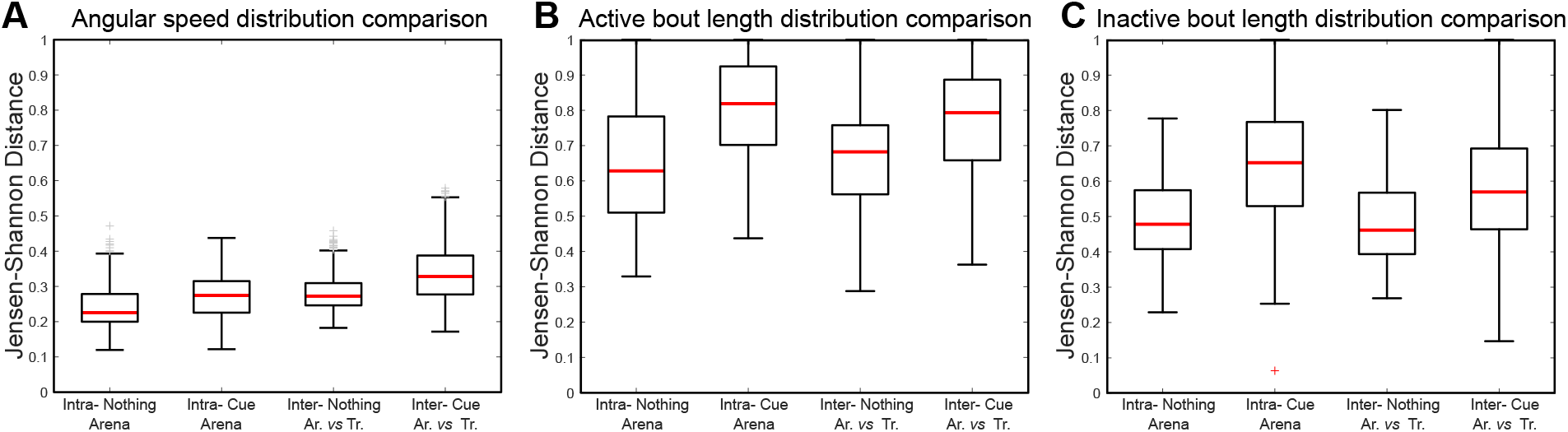
Comparison of angular speed and active-inactive bout durations distribution by Jensen-Shannon distance between arena and treadmill experiments. (A) Jensen-Shannon distance of angular speed distribution between trials in the arena and on the treadmill (Inter-), for both uniform and cue condition, are compared with the distance measured in-between different arena trials of both condition (Intra-). There is a significant impact of the treadmill and of the condition, as well as an interaction of both, on the distribution of the angular speed observed (Anova treadmill effect: *F* = 210.62, *p* < 0.0001; Cue effect: *F* = 132.4, *p* < 0.0001; Interaction: *F* = 17.38, *p* < 0.0001). However, in term of magnitude the influance of the treadmill is small: JS mean distance for Intra-groups is around 0.2 to 0.3 and around 0.3 and 0.35 for Inter-groups. (B) Jensen-Shannon distance of active bout duration distribution between trials in the arena and on the treadmill (Inter-), for both uniform and cue condition, are compared with the distance measured in-between different arena trials of both condition (Intra-). Anova treadmill effect: *F* = 1.75, *p* = 0.1859; Cue effect: *F* = 289.09, *p* < 0.0001; Interaction: *F* = 8.14, *p* = 0.0044. (C) Jensen-Shannon distance of inactive bout duration between trials in the arena and on the treadmill (Inter-), for both uniform and cue condition, are compared with the distance measured in-between different arena trials of both condition (Intra-). Anova treadmill effect: *F* = 29.76, *p* < 0.0001; Cue effect: *F* = 325.57, *p* < 0.0001; Interaction: *F* = 20.24, *p* < 0.0001.

## 3 Relative speed analysis

As we did not track the motion of the ball we could not extract the absolute speed of the ant on the treadmill. Nevertheless, we can infer a relative speed approximation from the distance to the center of the servosphere. To keep this distinction, and because of the PD, we kept the speed index on the treadmill expressed in pixels.s^−1^ rather than converting in mm.s^−1^. We therefore evaluated the relationship between ants on the treadmill with the one observed in the arena, to estimate the conservation of the motion pattern (figure A.3).

**Figure A.3:**
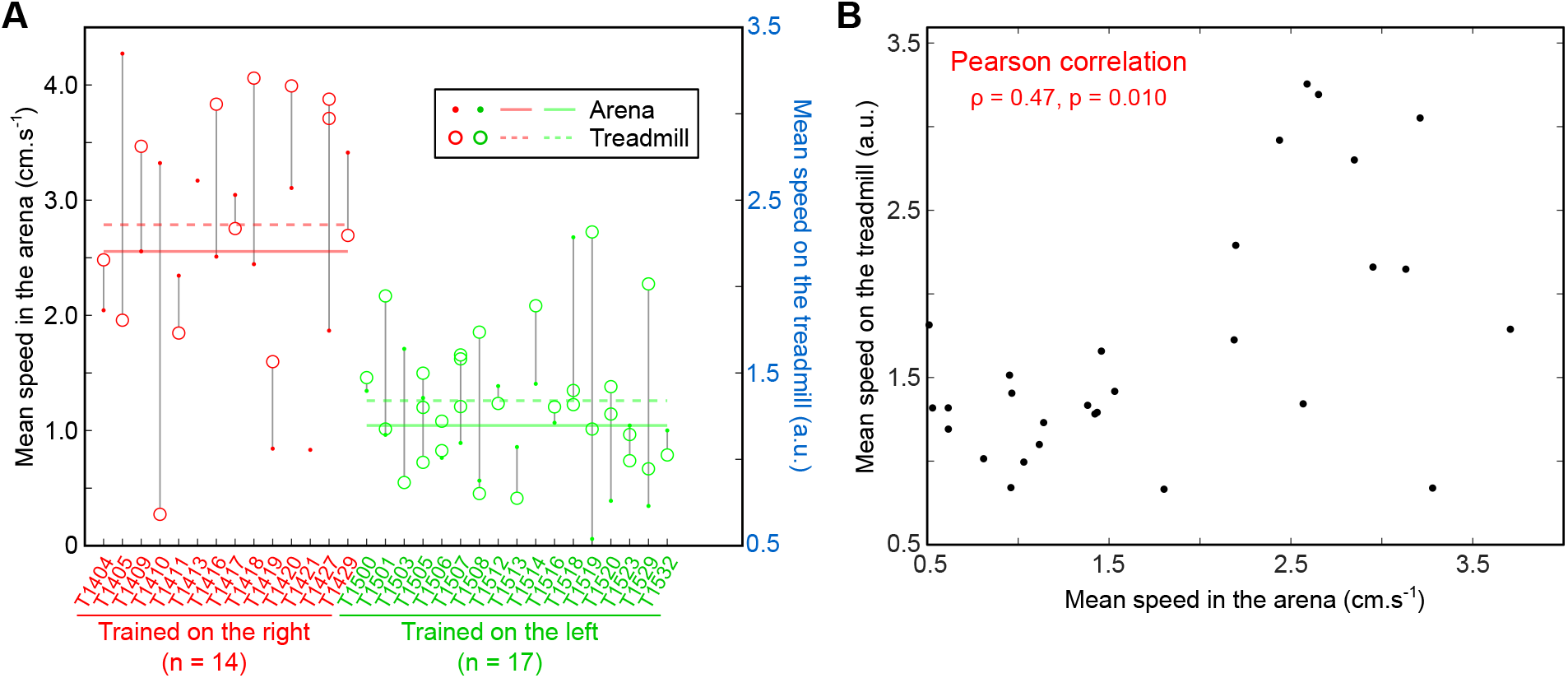
Correlation between individual mean speed observed during the arena training (last session) and on the treadmill. (A) For each individual, mean speed observed in the arena during the last session of training (colored points, left scale) and on the treadmill (colored circle, right scale). We observed a clear difference in term of the mean speed expressed by individuals during the training session with the feeder on the right (red) or on the left (green). This difference appeared also during the test on the treadmill, suggesting the preservation of individual traits of motion. The plain lines indicate the global mean for arena experiments and the dotted lines on the treadmill. (B) Correlation between mean speed observed in the arena during the last session of training and on the treadmill.

